# Diversity and modularity of tyrosine-accepting tRNA-like structures

**DOI:** 10.1101/2023.07.08.548219

**Authors:** Madeline E. Sherlock, Conner J. Langeberg, Jeffrey S. Kieft

## Abstract

Certain positive-sense single-stranded RNA viruses contain elements at their 3’ termini that structurally mimic tRNAs. These tRNA-like structures (TLSs) are classified based on which amino acid is covalently added to the 3’ end by host aminoacyl-tRNA synthetase. Recently, a cryoEM reconstruction of a representative tyrosine-accepting tRNA-like structure (TLS^Tyr^) from brome mosaic virus (BMV) revealed a unique mode of recognition of the viral anticodon-mimicking domain by tyrosyl-tRNA synthetase. Some viruses in the *hordeivirus* genus of *Virgaviridae* are also selectively aminoacylated with tyrosine, yet these TLS RNAs have a different architecture in the 5’ domain that comprises the atypical anticodon loop mimic. Herein, we present bioinformatic and biochemical data supporting a distinct secondary structure for the 5′ domain of the *hordeivirus* TLS^Tyr^ compared to those in *Bromoviridae*. Despite forming a different secondary structure, the 5′ domain is necessary to achieve robust *in vitro* aminoacylation. Furthermore, a chimeric RNA containing the 5′ domain from the BMV TLS^Tyr^ and the 3′ domain from a *hordeivirus* TLS^Tyr^ are aminoacylated, illustrating modularity in these structured RNA elements. We propose that the structurally distinct 5′ domain of the *hordeivirus* TLS^Tyr^s performs the same role in mimicking the anticodon loop as its counterpart in the BMV TLS^Tyr^. Finally, these structurally and phylogenetically divergent types of TLS^Tyr^ provide insight into the evolutionary connections between all classes of viral tRNA-like structures.

## INTRODUCTION

Viruses have evolved many strategies to promote translation and replication of their genome while evading host cell anti-viral surveillance pathways. In many cases, specifically-structured RNA elements in the viral genome’s non-coding regions play critical roles during infection, such as promoting viral protein synthesis and preventing viral RNA from degradation (Dreher 2010; Moon et al. 2012; Jaafar and Kieft 2019). Examples of multifunctional viral RNA elements are the tRNA-like structures (TLS) in many positive-sense, single-stranded, plant-infecting viruses (Hall 1979; Dreher 2010), and a few insect-infecting viruses (Gordon et al. 1995; Sherlock et al. 2021). These RNA elements are located at the termini of the 3’ UTR within the viral RNA and have been implicated in enhancement of translation, protection from 3’ to 5’ degradation, viral replication, and viral packaging (Joshi et al. 1985; Dreher and Hall 1988; Rao et al. 1989; Mans et al. 1992; Osman et al. 2000; Matsuda and Dreher 2004; Dreher 2009; Rao and Kao 2015).

As their name suggests, TLSs contain features that structurally mimic canonical tRNA, enabling interactions with certain host factors that normally process cellular tRNAs (Litvak et al. 1973; Rietveld et al. 1984; Goodwin and Dreher 1998; Hammond et al. 2009; Colussi et al. 2014; Bonilla et al. 2021). Notably, viruses that contain an authentic TLS have a canonical tRNA ‘CCA’ tail appended to the 3’ end of a pseudoknot that mimics the acceptor stem, and they undergo aminoacylation by a host amino-acyl tRNA synthetase (AARS) (Rietveld et al. 1984; Pleij et al. 1985; Dreher and Hall 1988; Bonilla et al. 2021; Langeberg et al. 2021; Sherlock et al. 2021).

TLSs are classified by the amino acid they are charged with, which correlates with sequence and structural features unique to each of the three classes. Valine-accepting tRNA-like structures (TLS^Val^) are found in the *Tymoviridae, Virgaviridae*, and *Alphatetraviridae* families and most closely resemble canonical tRNA with elements mimicking the D, T, and anticodon stems of tRNA^Val^ (Giegé et al. 1978; Dreher et al. 1992; Colussi et al. 2014; Hartwick et al. 2018; Sherlock et al. 2021). Histidine-accepting tRNA-like structures (TLS^His^) are found in *Virgaviridae* and *Tymoviridae* and retain some canonical tRNA features (D-loop/T-loop mimic) but are larger than tRNAs and contain certain secondary structure elements that are dissimilar to tRNA, including a long anticodon stem mimic with a large asymmetric internal loop (Rietveld et al. 1984; Joshi et al. 1985; Langeberg et al. 2021). Tyrosine-accepting tRNA-like structures (TLS^Tyr^) are found in *Virgaviridae* and *Bromoviridae*, contain even fewer features of canonical tRNA, and are much larger in size than tRNAs (Loesch-Fries and Hall 1982; Dreher and Hall 1988; Felden et al. 1994; Bonilla et al. 2021).

Recently, structures of the TLS^Tyr^ from brome mosaic virus (BMV) in both the unbound (RNA-only) and bound (RNA-AARS complex) were determined using single-particle cryo-electron microscopy (cryoEM) (Bonilla et al. 2021). Models of the BMV TLS resulting from these cryoEM reconstructions revealed the overall architecture of the seven helical domains of the RNA (Fig. 1A) as well as an unexpected binding mode with the host tyrosyl-tRNA synthetase (TyrRS) involving the anticodon recognition domain. Specifically, in the unbound form, the BMV TLS anticodon-mimicking domain is mobile, sampling a variety of positions and is thus not pre-organized to interact with the TyrRS (Bonilla et al. 2021). However, in the BMV TLS-TryRS complex, this domain is stably placed and oriented such that it is nearly parallel with the acceptor stem, contrasting with the perpendicular orientation resulting from the L-shape of tRNA.

**FIGURE 1.**
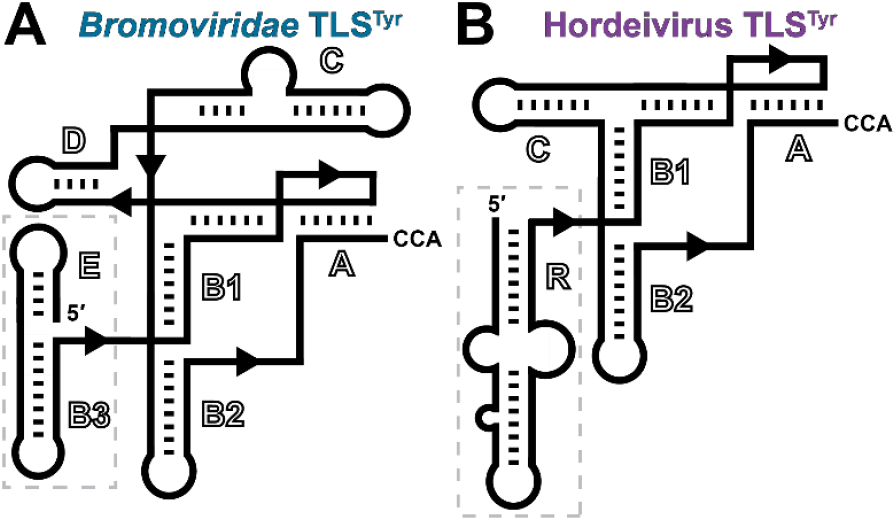
Cartoon diagrams of the secondary structure of the tyrosine-accepting tRNA-like structures (TLS^Tyr^) from the (*A*) *anulavirus, bromovirus, cucumovirus*, and *oleavirus* genera of *Bromoviridae* and (*B*) *hordeivirus* genus of *Virgaviridae*. Helical elements are lettered according to previous literature (Ahlquist et al. 1981; Solovyev et al. 1996; Savenkov et al. 1998; Bonilla et al. 2021). The respective 5’ domains, consisting of the (*A*) E and B3 stem loops or (*B*) R stem loop, are boxed in gray. The secondary structure of the R domain of the *hordeivirus* TLS^Tyr^ was not experimentally probed before this study, the putative structure is shown.

As mentioned above, TLS^Tyr^ are also found in viruses in the *Virgaviridae* family, and these TLS^Tyr^ representatives contain key differences compared to the BMV TLS. Specifically, all three members of the *hordeivirus* genus within *Virgaviridae* – barley stripe mosaic virus (BSMV), poa semilatent virus (PSLV), and lychnis ringspot virus (LRSV) – contain TLS^Tyr^s in their 3’ UTRs (Solovyev et al. 1996). The most notable difference between the *hordeivirus* TLS^Tyr^s and the BMV TLS (and related structures in other members of *Bromoviridae*) is that the former do not contain the E and B3 stem loops (Fig. 1A). This is puzzling as these elements function as the BMV TLS’s anticodon-mimicking domain (Felden et al. 1994; Bonilla et al. 2021). Instead, the 5’ domain for *hordeivirus* TLSs was proposed (Kozlov et al. 1984; Solovyev et al. 1996; Savenkov et al. 1998) to comprise a large stem-loop structure with an asymmetric internal loop, labeled ‘R’ (Fig. 1B). Despite these differences, the aminoacylation of the BSMV genome with tyrosine at its 3’ terminus was demonstrated over four decades ago (Agranovsky et al. 1981; Agranovsky et al. 1982; Loesch-Fries and Hall 1982). However, key features of these divergent TLS^Tyr^, such as the minimal tyrosylation-competent element and therefore the necessary domains within the 3’ UTR that comprise the TLS, have not been experimentally determined. Also, while secondary structure models for the 3’ UTR, including the putative TLS domain, have been proposed for all three members of the *hordeivirus* genus (Kozlov et al. 1984; Agranovsky et al. 1992; Solovyev et al. 1996; Savenkov et al. 1998), these have not been probed experimentally.

The discrepancy as to the nature of a putative anticodon-mimicking domain in the TLS^Tyr^ representatives in *hordeivirus* as well as other subtle differences compared to TLS^Tyr^s in *Bromoviridae*, motivated us to experimentally interrogate the structural features of these RNAs. Chemical probing data combined with functional assays enabled us to compare domains of *hordeivirus* TLS^Tyr^s with those of BMV in the context of its known structure. We have assigned the role of anticodon mimicry to the 5’ domain of *hordeivirus* TLS^Tyr^, which is required for tyrosylation and much larger than its counterpart in BMV. We then identified the most highly conserved features and areas of divergence among all TLS^Tyr^ examples to illuminate the most critical features for aminoacylation function across this diverse TLS class.

## RESULTS AND DISCUSSION

### Experimental exploration of *hordeivirus* TLS^Tyr^s secondary structures

We first experimentally interrogated the proposed *hordeivirus* TLS^Tyr^s secondary structures compared to *Bromoviridae* TLS^Tyr^s. Selective 2’ hydroxyl acylation by primer extension (SHAPE) probing was used to evaluate the previously proposed structures by mapping reactivity of the chemical modifier for each RNA onto its secondary structure model. Overall, the chemical probing data for all three *hordeivirus* TLSs (Figs. 2A, S1) are consistent with previously proposed secondary structure predictions based solely on sequence information (Kozlov et al. 1984; Agranovsky et al. 1992; Solovyev et al. 1996). Notably, the 5’ portion of all three *hordeivirus* TLS^Tyr^ folds into a long stem-loop containing an asymmetric internal loop and a single bulged nucleotide; hereafter we refer to the 5’ region as the R domain. The remaining portion of the *hordeivirus* TLSs^Tyr^ folds in a manner that is largely consistent with *Bromoviridae* TLS^Tyr^ RNAs, with the base-paired regions in A, B1, B2, and C all displaying low reactivity to the chemical modifier. Notably, the pyrimidine-rich linker between B2 and the closing of the acceptor stem pseudoknot (A) shows low reactivity in all three *hordeivirus* TLS^Tyr^s. The pyrimidine-rich linker region in the BMV TLS^Tyr^ also demonstrates low chemical probing reactivity, likely due to a stable conformation against the B1 stem and the loop of the E stem as seen in the cryoEM structure (Bonilla et al. 2021).

**FIGURE 2.**
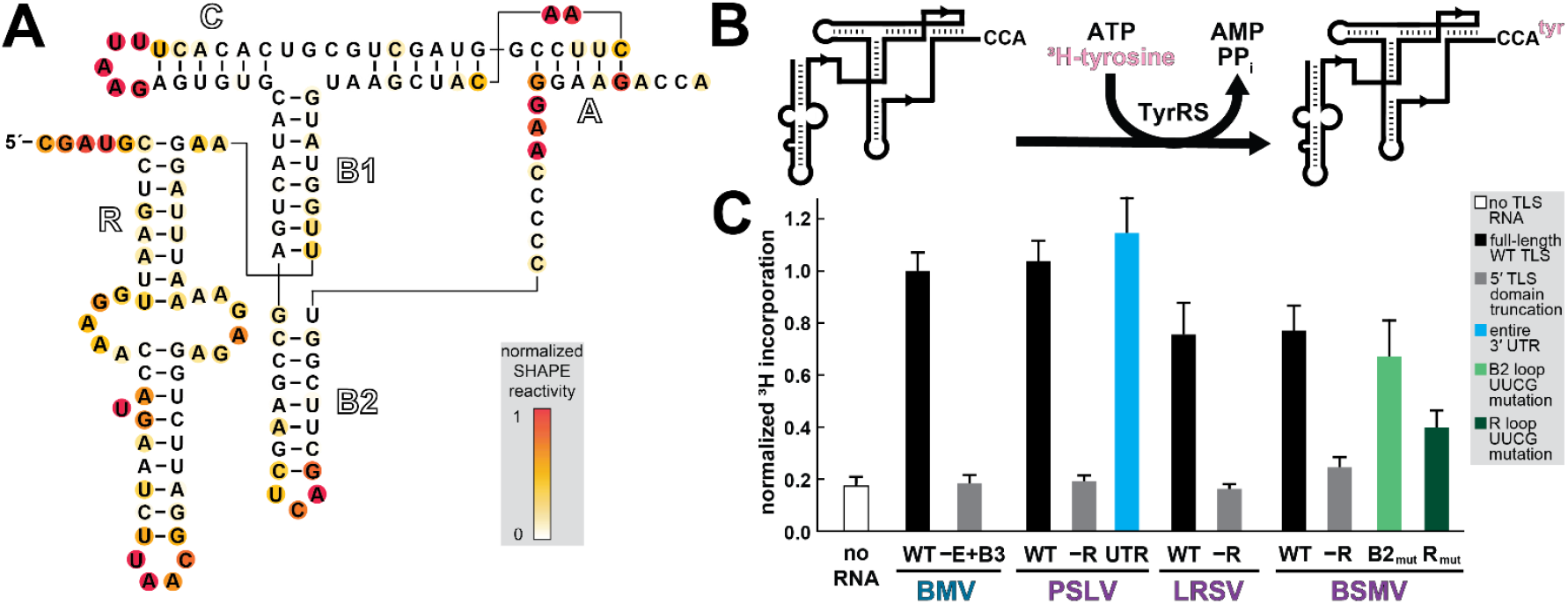
Structure and function of *hordeivirus* TLS^Tyr^. (*A*) Chemical probing of the TLS representative from BSMV RNA1 using the SHAPE reagent NMIA. Reactivity was background subtracted and normalized according to the reactivity of loop regions in hairpin structures (not shown) flanking the TLS structure on both the 5′ and 3′ ends. See Table S1 for complete sequence details and Fig. S1 for chemical probing of additional hordeivirus TLS^Tyr^ RNAs. (*B*) Schematic of *in vitro* aminoacylation assay using *in vitro* transcribed RNA, recombinantly-expressed and purified tyrosyl-tRNA synthetase (TyrRS), and ^3^H-labeled tyrosine. (*C*) Relative activity of TyrRS on various TLS^Tyr^ RNAs as measured by the covalent addition of radiolabeled ^3^H-tyrosine at their 3′ termini. ^3^H incorporation was normalized to the WT BMV TLS^Tyr^ construct, which had been previously tested (Bonilla et al. 2021). The truncated constructs (-E+B3 or -R) are missing their respective 5’ domains, as indicated by the dashed box in Fig. 1, and begin four nucleotides prior to the B1 stem. The PSLV full 3’ UTR RNA includes the entire sequence after the stop codon. The BSMV TLS RNAs with mutations to the B2 and R stem loops (B2_mut_ and R_mut_) contain UUCG tetraloops in place of the native loop sequence (see panel *A* for the WT BSMV TLS^Tyr^ sequence).

### Defining the minimal functional unit for the *hordeivirus* TLS^Tyr^

We next determined which portions of *hordeivirus* 3’ UTR comprise the minimal unit competent for tyrosylation, as previous studies used the entirety of the 3’ UTR as a substrate (Agranovsky et al. 1982; Agranovsky et al. 1992). We used an *in vitro* assay with recombinantly expressed and purified TyrRS previously shown to be active on the BMV TLS^Tyr^ in a dose-dependent manner, and inactive on a TLS^Val^ (Bonilla et al. 2021). In this assay, aminoacylation with ^3^H-tyrosine leads to ^3^H-labeled RNA (Fig. 2B), which is quantified and normalized to the wild-type (WT) full-length BMV TLS^Tyr^. Validating the assay, there was no measurable tyrosylation above the no RNA (negative) control for a BMV TLS^Tyr^ that is truncated to lack the E and B3 stem loops (Fig. 2B), consistent with previous data showing the BMV TLS^Tyr^ is sensitive to mutations in the B3 loop (Bonilla et al. 2021).

We then tested three putative *hordeivirus* RNAs, each beginning five nucleotides prior to the R domain. All were tyrosylated to a similar level to the BMV TLS^Tyr^. Truncations to all three *hordeivirus* TLS^Tyr^s that removed the R domain decreased tyrosylation to levels similar to the negative control. Thus, the R domain is critical for aminoacylation of the *hordeivirus* TLS^Tyr^s, perhaps analogous to the BMV TLS^Tyr^ 5’ domain comprising the E and B3 stem-loops, referred to hereafter as ‘E+B3’. Furthermore, an RNA containing the entire 3’ UTR of PSLV is tyrosylated to a similar level as the PSLV TLS^Tyr^ that begins just prior to the R domain (Fig. 2C). These results demonstrate that the R domain is a required functional component for the *hordeivirus* TLS^Tyr^s, but other elements in the 3’ UTR upstream of the R domain are not necessary for aminoacylation (discussed in greater detail in the Concluding Remarks).

To further test the specific involvement of the *hordeivirus* TLS^Tyr^ R stem in tyrosylation, we mutated its apical loop to a UUCG tetraloop. This mutation substantially decreased tyrosylation, further illustrating the importance of this element. In contrast, mutation of the *hordeivirus* TLS^Tyr^ B2 apical loop had a less deleterious effect on tyrosylation (Fig. 2C). These results are consistent with analogous mutations to the BMV TLS B3 and B2 apical loops, respectively (Bonilla et al. 2021).

### Anticodon mimicry by the 5’ domain of *hordeivirus* TLS^Tyr^

After confirming the secondary structure of the R domain (Fig. 2A) and determining that this region of the *hordeivirus* TLS^Tyr^ is a necessary component for aminoacylation (Fig. 2C), we next investigated whether the R domain participates in anticodon mimicry analogously to the B3 domain of BMV TLS (Felden et al. 1994; Bonilla et al. 2021). In the R domain, the putative anticodon-mimicking loop for BSMV (i.e. the apical loop) is an ‘UAAC’ tetraloop, which differs slightly from the BMV B3 apical loop (UACA). Neither match the canonical tRNA^Tyr^ GUA anticodon embedded in a seven-nucleotide loop. Nonetheless, mutation of either the BSMV R domain or BMV TLS^Tyr^ B3 apical loops to UUCG tetraloops resulted in the loss of aminoacylation (Fig. 2C) (Bonilla et al. 2021). Additionally, the R domain of *hordeivirus* TLS^Tyr^s contains a much longer helical region (18 total base pairs) than the combined length of the E+B3 domains of BMV (12 total base pairs).

Despite these differences, we hypothesized that the R domain of the *hordeivirus* TLSs serves the same functional role as the E+B3 domain of BMV, including interactions with the anticodon-recognition domain of TyrRS. If so, they could be functionally interchangeable. To test this, we created chimeric BMV and BSMV TLS^Tyr^ RNAs by appending the E+B3 domain of BMV to the BSMV TLS (Fig. 3A), and the R domain of BSMV to the BMV TLS (Fig. 3B). We then tested whether these chimeric constructs were competent substrates for tyrosylation *in vitro*.

**FIGURE 3.**
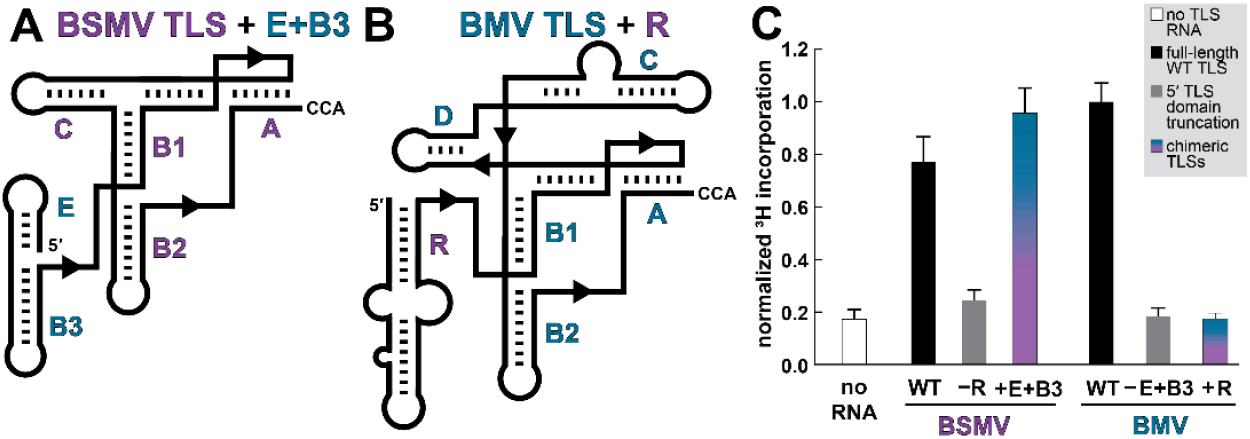
Aminoacylation of chimeric TLS^Tyr^s. (*A,B*) Cartoon diagrams of the secondary structure of the TLS^Tyr^ from (*A*) BSMV with the E and B3 stem-loops of BMV appended to the 5’ end in lieu of the R domain and (*B*) BMV with the R stem-loop of BSMV appended to the 5’ end in lieu of the E+B3 domain. (*C*) Relative activity of TyrRS on WT, truncated (-R or -E+B3, gray), and chimeric (+E+B3 or +R, blue-purple) TLS^Tyr^ RNAs as measured by ^3^H-tyrosine incorporation. See legend to Fig. 2 for additional details.

Again, the 5’ truncated BSMV TLS^Tyr^ without the R domain has no detectable aminoacylation activity above the no RNA negative control (Fig. 2C). However, the addition of the E+B3 domain to the BSMV *hordeivirus* TLS^Tyr^ in place of the R domain fully restores tyrosylatability (Fig. 3C). Thus, loss of function due to removal of the R domain is restored by adding the known anticodon-mimicking domain of the BMV TLS. Despite differences in size, primary sequence, and architecture, the R and E+B3 domains perform homologous functions. This could be accomplished by forming a similar structural architecture with the apical loop of the R domain contacting the anticodon recognition domain of the TyrRS enzyme similarly to the apical B3 loop. We speculate that the internal loop nucleotides in the R domain could allow it to be structurally flexible to contact the anticodon recognition domain of TyrRS, perhaps in a parallel conformation as observed for the BMV TLS^Tyr^ (Bonilla et al. 2021).

Interestingly, the other chimeric construct with the R domain of the BSMV appended to the 3’ portion of the BMV TLS^Tyr^ (Fig. 3B) did not restore aminoacylation activity (Fig. 3C). Chemical probing data show that this chimeric RNA is likely not globally misfolded (Fig. S2). Rather, we hypothesize that the much larger R domain compared to the size of E+B3 is somehow incompatible with the overall structure of the BMV TLS^Tyr^. Some possible explanations are that the R domain is incompatible with a TLS^Tyr^ containing both the C and D stem-loops. Or, in the BMV TLS^Tyr^ there is necessary contact between E+B3 and B2 that cannot be formed by the larger R stem-loop in the context of BMV. However, in the absence of a high-resolution structure of a *hordeivirus* TLS^Tyr^ RNA, it is difficult to determine whether steric clashes, the absence of critical tertiary contacts, or other factors lead to the inability of the chimeric BMV TLS^Tyr^ + BSMV R construct to be aminoacylated.

### Unifying and distinguishing features between all viral tyrosine TLSs

To evaluate the conservation and variations across all known TLS^Tyr^s, we created sequence and secondary structure alignments using 32 unique TLS^Tyr^ sequences from 13 distinct viruses in *Bromoviridae*, and the eight unique TLS^Tyr^ sequences from the three members of *hordeivirus* (Supplemental Files 1 and 2). We used R2R (Weinberg and Breaker 2011) to depict the consensus sequence and secondary structure models (Fig. 4A,B).

**FIGURE 4.**
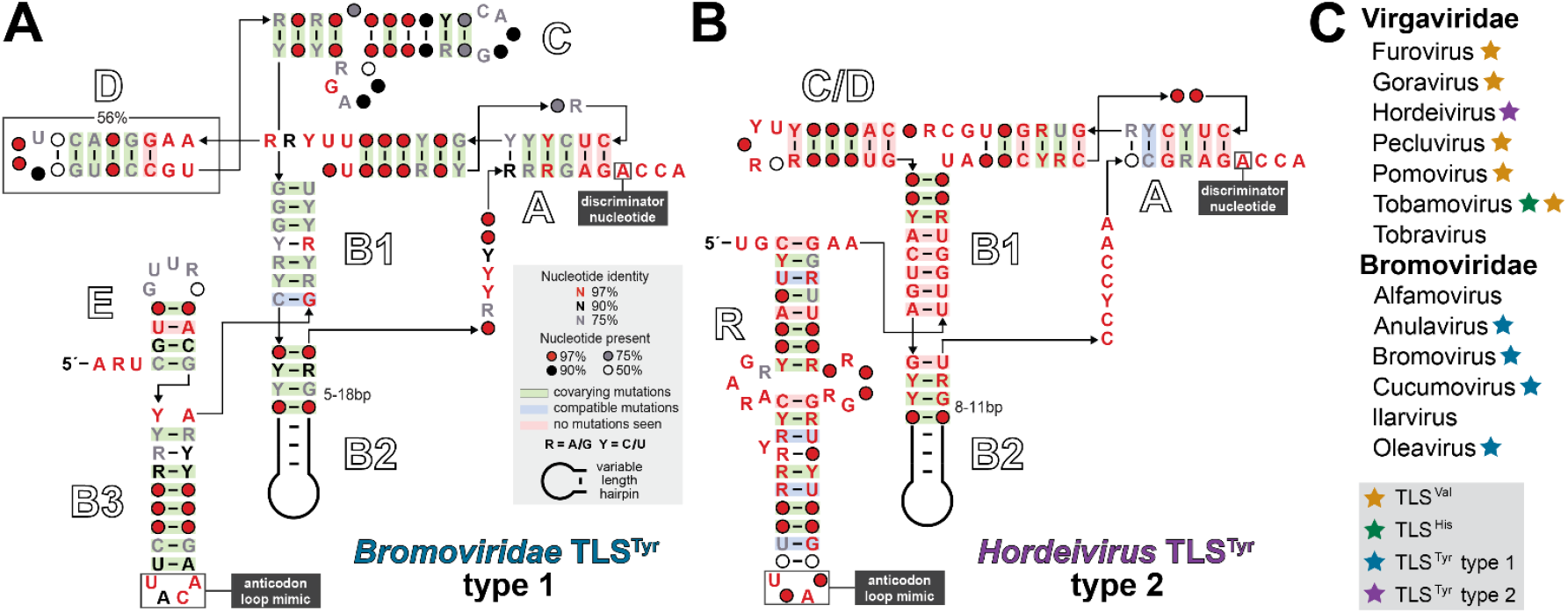
Conservation and phylogenetic distribution of viral tyrosine-accepting tRNA-like structures. (*A*) Consensus sequence and secondary structure model for the 32 unique TLS^Tyr^ sequences from 13 distinct viruses in *Bromoviridae*. (*B*) Consensus sequence and secondary structure model for the 8 unique TLS^Tyr^ sequences from 3 distinct viruses in *Virgaviridae*. The stem loop labeled as C/D was referred to as the C stem in previous literature; however, it may more closely resemble the optional D stem found in type 1 TLS^Tyr^. (*C*) Phylogenetic distribution of all types of TLSs in the *Virgaviridae* and *Bromoviridae* families.

All TLS^Tyr^s contain a highly conserved 3’ end; the identity of the discriminator base – the nucleotide immediately preceding the terminal CCA – is perfectly conserved as an adenosine, and the terminal base pair of the acceptor stem-mimicking pseudoknot is also perfectly conserved as a C-G. This high degree of conservation is perhaps unsurprising, given that the adenosine discriminator nucleotide and the C-G pair at the base of the acceptor stem are the most critical identity elements for recognition of canonical tRNA^Tyr^ by TyrRS (Fechter et al. 2000; Fechter et al. 2001; Tsunoda et al. 2007). While the R and E+B3 domains diverge between the two subclasses of TLS^Tyr^, both contain tetranucleotide apical loops that typically start with a U followed by three A or C nucleotides. Again, this sequence differs from the tRNA^Tyr^ anticodon sequence, but in tRNA^Tyr^ the anticodon is a weaker identity element for TyrRS recognition compared to the acceptor stem and discriminator nucleotide (Fechter et al. 2001). Specifically, most mutations to the tRNA^Tyr^ anticodon result in <10-fold loss in aminoacylation efficiency while mutations to the acceptor stem and discriminator nucleotide result in a >1000-fold decrease or complete loss in aminoacylation (Fechter et al. 2001). Although the identity of the nucleotides in the anticodon mimicking loop (B3 for BMV; R for BSMV) differ from canonical tRNA^Tyr^, mutations to the loop sequence for TLS^Tyr^ decreases aminoacylation efficiency (Fig. 2C) (Bonilla et al. 2021).

An area of variability among TLS^Tyr^ is in the region located between the B1 stem and the A pseudoknot. In the BMV TLS^Tyr^, this region comprises the C and D stem-loops, which are not universally conserved. The C stem-loop is always present in *Bromoviridae* TLS^Tyr^s and contains elements critical for recognition by the RNA-dependent RNA polymerase (RdRP) to achieve viral replication (Rao et al. 1989; Chapman and Kao 1999; Rao and Kao 2015). The cryoEM structure of the BMV TLS^Tyr^ revealed that the C stem-loop is close in space to the A pseudoknot (Bonilla et al. 2021), consistent with its importance for replication as transcription begins at the 3’ end. The D stem-loop is not always present for *Bromoviridae* TLS^Tyr^s (Ahlquist et al. 1981; Bonilla et al. 2021); when present, its function is unknown.

The *hordeivirus* TLS^Tyr^s have a single stem-loop in in this region, which we have labeled C throughout in accordance with previous literature, and are proposed to be missing a D stem-loop (Savenkov et al. 1998). However, the *hordeivirus* TLS^Tyr^s C stem-loop differs significantly from the C stem-loop of *Bromoviridae* TLSs. First, the length of the helix in hordeiviruses (6bp with a pentanucleotide apical loop) is dissimilar to the *Bromoviridae* TLS^Tyr^ C stem (10 bp with a large, asymmetric internal loop). Second, the proximity of the C stem to the 3’ end to perform its role in replication might not be relevant for the *hordeivirus* TLS^Tyr^. As members of the *Virgaviridae* family, these viruses have a different RdRP lineage compared to *Bromoviridae*. The *hordeivirus* TLS^Tyr^s might not rely on this additional C stem loop, but instead contain recognition requirements for replication more consistent with the other *Virgaviridae* TLS classes (Singh and Dreher 1997; Deiman et al. 1998; Osman et al. 2000). We propose that the stem loop in this region in *hordeivirus* TLSs might have more similarity to the BMV D stem-loop and have labeled it ‘C/D’ (Fig. 4B) to note this ambiguity.

### TLS^Tyr^s differ in the structures directly upstream

An intriguing connection between the viruses within *hordeivirus* and other genera within *Virgaviridae* (Fig. 4C) is the overall architecture of the entire 3’ UTR upstream of the TLS domain. The TLS^His^ from tobacco mosaic virus (TMV) is preceded by an upstream pseudoknot domain (UPD) consisting of three consecutive pseudoknots connected by linker and loop nucleotides (Van Belkum et al. 1985; Zeenko et al. 2002). For TMV, a portion of the UPD – two of the pseudoknots and the intervening loop nucleotides – has been implicated as necessary for enhancing translation and is the binding site for elongation factor eEF1A (Zeenko et al. 2002). The translation enhancement effect of the UPD was increased synergistically with the 5’ cap structure and elements in the 5’ UTR (Leathers et al. 1993). Both BSMV and PSLV contain the same UPD architecture as TMV, including the highly conserved loop nucleotides and three pseudoknots (Van Belkum et al. 1985; Zeenko et al. 2002). Chemical probing data support the predicted secondary structure of this region for both BSMV and PSLV, and the presence of the UPD does not appear to affect folding of the TLS (Fig. S3) or be necessary for aminoacylation (Fig. 2C).

An additional layer of complexity in the *hordeivirus* 3’ UTRs are poly(A) sequences. Both BSMV (contains a UPD) and LRSV (no UPD) contain a poly(A) stretch of variable length (4-40 nt) directly after the stop codon (Agranovsky et al. 1982; Jackson et al. 1989; Agranovsky et al. 1992; Solovyev et al. 1996), which is not required for replication (Zhou and Jackson 1996). In contrast, PSLV (contains a UPD) has several short (4-5 nt), interspersed poly(A) stretches between the stop codon and UPD (Agranovsky et al. 1992). It has been proposed (Dreher and Miller 2006) that 3’ UTR elements in other viruses within *Virgaviridae* might contribute to 5’ to 3’ end communication similarly to eukaryotic mRNAs – a role that poly(A) sequences in *hordeivirus* genomes could provide through poly(A) binding protein. However, the relative contributions of the TLS, UPD, and poly(A) features to viral translation enhancement has not been investigated in *hordeivirus* to date, and much remains to be unraveled in the function of these multi-functional, highly-structured, non-coding RNA elements.

### Concluding Remarks

The data presented herein define the functional element necessary to achieve aminoacylation for *hordeivirus* tyrosine-accepting TLSs. We have shown that certain portions of TLS^Tyr^s are similar among all representatives, with the highest degree of conservation at the extreme 3’ end. Other portions of the TLS^Tyr^ are much more variable and some features may be more relevant to functions other than aminoacylation. The 5’ domains of the two TLS^Tyr^ types appear dissimilar in terms of secondary structure, but our mutational analyses suggest a common function, which also demonstrates the modularity of RNA structure and function.

While the evolutionary trajectory of all TLS-containing viruses is difficult to definitively determine, it has been proposed that these RNAs originated in the *Virgaviridae* family, since all three known TLS classes are represented in this one viral family (Fig. 1A) (Dreher 2010). While the amino acid specificity of the *hordeivirus* TLS class for tyrosine matches that of *Bromoviridae*, other features of the *hordeivirus* genus align more closely with other members of *Virgaviridae*, including the same UPD in some members of both *hordeivirus* and *tobamovirus*. The *hordeivirus* TLS R domain, which we propose functions as an anticodon mimic, has some intriguing similarities to the anticodon stem mimic of TLS^His^: both have similar overall lengths, have an asymmetric internal loop, and the trinucleotide anticodon mimic contained in the apical loop is divergent from the 7-nucleotide canonical tRNA anticodon loop. It is tempting to speculate that the original tyrosine-accepting TLSs could have emerged in *hordeivirus*, having diverged at some point from the histidine class in *tobamovirus*. These TLS^Tyr^ could have subsequently appeared in *Bromoviridae*, possibly through recombination. Regardless of the evolutionary trajectory, the presence of different TLS classes with other RNA elements in the 3’ UTR represent an intriguing set of RNA structures contributing to numerous functions in the viral lifecycle. Despite recent advances in understanding the structural tRNA mimicry of these TLS domains, the individual roles and potential synergistic effects of these viral RNA structures in translation, replication, and packaging remain largely mysterious.

## MATERIALS AND METHODS

### RNA consensus sequence and secondary structure model

An existing alignment (Rfam ID: RF01075) of two *hordeivirus* TLSs (BSMV RNA1 and PSLV RNA1) was obtained from the Rfam database version 13.0 (Kalvari et al. 2018). This preliminary alignment was extended on the 5’ end to include the R domain. This domain was previously excluded as it had not been confirmed as necessary for function. Sequences of additional examples of this class (BSMV RNA2 & RNA3, PSLV RNA2, LRSV RNA1 & RNA2) were retrieved from the National Center for Biotechnology Information (NCBI) viral nucleotide database (https://www.ncbi.nlm.nih.gov/labs/virus/vssi/#/), then manually added and aligned. A previously published alignment of *Bromoviridae* TLSs (Bonilla et al. 2021) was amended to only include one unique example per viral RNA (previously all 512 unique sequences were reported, which included many examples from different strains of the same virus) to be consistent with the alignment of *hordeivirus* TLSs, as described above. Additionally, the sequences from tomato aspermy virus, which is not tyrosylated (Joshi and Haenni 1986) and therefore does not functionally belong to this class despite sequence and secondary structure similarity, were removed from the *Bromoviridae* TLS alignment. The consensus sequence and secondary structure models for the *hordeivirus* and *Bromoviridae* TLS^Tyr^ classes were separately calculated and visualized, using the two above alignments (*hordeivirus*: Supplemental File 1; *Bromoviridae*: Supplemental File 2) as input files, using R2R (Weinberg and Breaker 2011) and labeled in Adobe Illustrator.

### RNA preparation

DNA templates were ordered as gBlock double-stranded DNA fragments (IDT) amplified by PCR in 100 μL volumes (see Supplementary Table S1 for all DNA template and primer sequences used in this study). Reverse primers used to amplify the constructs for in vitro aminoacylation assays contained two 5′-terminal 2′OMe-modified bases to achieve a higher yield with the correct 3′ end of the construct. dsDNA amplification was confirmed by 1% agarose gel electrophoresis. Transcriptions were performed in 200 μL volume using 50 μL of PCR product as template dsDNA, 30 mM Tris pH 8.0, 60 mM MgCl_2_, 8 mM each NTP, 10 mM DTT, 0.1% spermidine, 0.1% Triton X-100, and T7 RNA polymerase. Reactions were incubated at 37°C overnight, then purified using denaturing 10% polyacrylamide gel electrophoresis (PAGE). Bands were visualized by UV shadowing, excised, and sliced into small pieces, and soaked in ∼300 μL of diethylpyrocarbonate (DEPC)-treated milli-Q filtered water (Millipore) containing 20 mM sodium acetate (pH 5.3) at 4°C overnight to elute the RNA. Supernatant-containing RNA was subjected to ethanol precipitation then resuspended in DEPC-treated water, diluted to the appropriate concentration, and stored at −20°C.

### Chemical probing of RNAs *in vitro*

Structure probing experiments using the chemical modifier *N*-methyl isatoic anhydride (NMIA) were performed as described previously (Cordero et al. 2014; Langeberg et al. 2021; Sherlock et al. 2021). Briefly, RNA was refolded by heating to 90°C for 5 min, cooled to 23°C, then incubated with 10 mM MgCl_2_ for 20 min. Subsequently, the refolded RNA was modified by incubating with NMIA for 15 min at 23°C. NMIA modification reaction conditions and final concentrations: 120 nM RNA, 6 mg/mL NMIA or dimethyl sulfoxide (DMSO; negative control), 50mM HEPES-KOH (pH 8.0), 10mM MgCl_2_, 3 nM 5-6 FAM-labeled RT primer (see Supplementary Table S1 for sequence). Modification was quenched by the addition of NaCl to 500 mM, Na-MES buffer (pH 6.0) to 50 mM, and oligo (dT) magnetic beads [Invitrogen Poly(A) PuristMAG Kit], which hybridize to the poly(A) stretch contained in the RT DNA primer.

Chemically-modified RNAs were recovered using oligo dT magnetic beads, washed with 70% ethanol, then resuspended in water. Reverse transcription was performed using SuperScript III enzyme (Invitrogen). RNA ladders were produced by four separate reverse transcription reactions using ddNTPs. Reverse transcription reactions were incubated at 50°C for 45 min, then the RNA was degraded by adding NaOH (final concentration 200 mM), heating to 90°C for 5 min, then quenching with an acidic solution (final concentration: 250 mM sodium acetate pH 5.2, 250 mM HCl, 500 mM NaCl). The remaining DNA products were purified using the magnetic stand then washed with 70% ethanol. A solution containing HiDi formamide solution (ThermoFisher) and spiked with GeneScan 500 ROX Dye Size Standard (ThermoFisher) was added to elute DNA products from the magnetic beads.

5′-FAM-labeled reverse-strand DNA products were analyzed by capillary electrophoresis using an Applied Biosystems 3500 XL instrument. Fragment size analysis, alignment, background subtraction, and normalization (based on reactivity in flanking stem–loop regions) were performed using the HiTrace RiboKit (https://ribokit.github.io/HiTRACE/) (Kladwang et al. 2011; Yoon et al. 2011; Kim et al. 2013; Lee et al. 2015) in MatLab (MathWorks), and figures were rendered using RiboPaint (https://ribokit.github.io/RiboPaint/) in MatLab, then labeled in Adobe Illustrator. SHAPE reactivity was superimposed on the previously proposed secondary structure models (Van Belkum et al. 1985; Solovyev et al. 1996).

### Aminoacylation assays

The TyrRS enzyme from a model host of BMV, *Phaseolus vulgaris* (common bean), used in the current study was recombinantly expressed and purified previously; details can be found in the Materials and Methods section of Bonilla et al. (2021). Aminoacylation constructs were refolded by heating to 90°C for 5 min, cooling to 23°C, then incubated with 10 mM MgCl_2_ for 20 min.

Aminoacylation reactions were set up by mixing 2 μL of 1 μM RNA or water, 1 μL of 10X aminoacylation buffer (10X: 300 mM HEPES-KOH [pH 7.5], 20 mM ATP, 300 mM KCl, 50 mM MgCl_2_, 50 mM DTT), 1 μL ^3^H-labeled L-tyrosine (60 Ci/mmol), 1 μL of 3 μM TyrRS enzyme (10X: 2 μM) and 5 μL of water (final reaction volume = 10 μL). Aminoacylation reactions, each performed in triplicate, were incubated at 30°C for 30 min, diluted by adding 200 μl of wash buffer (1X: 20 mM Bis-Tris [pH 6.5], 10 mM NaCl, 1 mM MgCl_2_, with trace xylene cyanol for visualization), then immediately loaded onto a vacuum filter Minifold I 96 well dotblot apparatus (Whatman). Using the dot-blot system, the reaction solution was filtered through a layer of Tuffryn membrane (PALL Life Sciences), Hybond positively charged membrane (GE Healthcare), and gel dryer filter paper (Bio-Rad). Prior to filter blotting apparatus assembly, each layer was equilibrated in 1X wash buffer without xylene cyanol. The RNA binds to the positively-charged Hybond membrane. After application of the reaction solution, each blot was immediately washed 3 times with 300 μL of wash buffer containing trace xylene cyanol. The filters were subsequently dried and the blots from the HyBond membrane were excised and measured for ^3^H incorporation by liquid scintillation counter (Perkin-Elmer Tri-Carb 2910 TR). Data were analyzed and plotted using Microsoft Excel, then labeled in Adobe Illustrator.

## SUPPLEMENTAL MATERIAL

Supplemental material is available for this article.

## AKNOWLEDGEMENTS

We thank Andrea MacFadden for assistance in purifying the TyrRS enzyme. We thank members of the Kieft laboratory for helpful discussions. This work was supported by National Institutes of Health/National Institute of General Medical Sciences (NIH/NIGMS) grant R35GM118070 to JSK. MES was a Jane Coffin Childs Postdoctoral Fellow.

